# Oligotrophy and organic carbon dissipation as trophic strategy in chernozem rare and uncultured taxa

**DOI:** 10.1101/2025.11.25.690425

**Authors:** K.A. Demin, V.V. Sedova, K.I. Krylov, P.A. Mirko, Y.N. Kocharovskaya, T.M. Minkina, E.V. Prazdnova

**Affiliations:** Southern Federal University, 344090, Rostov-on-Don, Russia; G.K. Skryabin Institute of Biochemistry and Physiology of Microorganisms, 142290, Pushchino, Moscow region, Russia

**Keywords:** Soil metagenomics, oligotrophy, rare biosphere, ecophysiology

## Abstract

Soil microbial communities harbor vast phylogenetic diversity, yet the functional ecology of rare and uncultured taxa remains poorly understood. These low-abundance microorganisms may employ specialized trophic strategies enabling their persistence in resource-limited environments. This study examined whether established oligotrophy markers could effectively characterize and differentiate the ecophysiology of rare and uncultured bacteria recovered from diverse Russian soil metagenomes. A total of 31 soil samples (chernozems, fluvisols, solonetz, solonchak, kastanozems, and leptosols) were collected from protected natural areas in the Rostov Region, Russia. Shotgun metagenomic sequencing was performed, and 246 metagenome-assembled genomes (MAGs) were recovered through assembly and binning. Genomic and functional traits associated with oligotrophy - including genome size, predicted generation time, ribosomal RNA operon copy number, two-component signaling systems, chemotaxis proteins, and carbohydrate-active enzymes - were analyzed. Metabolic capabilities for C1-compound oxidation, benzoyl-CoA pathway utilization, organosulfonate metabolism, and atmospheric trace gas scavenging were annotated. Principal component analysis was employed to cluster MAGs based on ecological strategy. MAGs spanned 18 bacterial phyla, with genome sizes ranging from 0.58–11.6 Mb and predicted doubling times from 0.6–15.6 hours. Six statistically significant clusters were identified, corresponding to distinct life-history strategies: fast-growing generalists, hydrogenotrophs, C1-compound specialists, polysaccharide degraders, aromatic compound degraders, and minimal-genome specialists. Oligotrophic lifestyles were confidently inferred for Methylomirabilota, Krumholzibacteriota, and Eisenbacteria MAGs, characterized by slow growth, reliance on low-molecular-weight carbon dissipation, and reduced regulatory complexity. Copiotrophic strategies were associated with Myxococcota, Bacteroidota, Gammaproteobacteria, and Verrucomicrobiota, which exhibited large genomes, rapid doubling times, extensive two-component systems, and high carbohydrate-active enzyme abundances. Combined analysis of genome size, generation time, regulatory system complexity, chemotaxis capacity, and substrate utilization pathways provides a robust framework for inferring trophic strategies of uncultured soil bacteria. Oligotrophy among rare taxa is characterized by adaptation to low-molecular-weight carbon dissipation, atmospheric trace gas oxidation, representing ecological strategies that enable persistence in nutrient-limited soil microhabitats.

## Introduction

The soil microbiome has been a central focus of ecological and biogeochemical research for decades. Soil microorganisms regulate terrestrial ecosystem functioning by mediating organic matter decomposition, transformation, and sequestration, and by driving global geochemical cycles (Kaviya et al., 2019). The structural and functional organization of microbial communities is, in turn, strongly modulated by edaphic properties. Soils represent intrinsically heterogeneous environments, especially with respect to the distribution and availability of nutrients. Spatial variations in resource availability result from geographic, geological, and climatic factors, combined with organic matter deposition and decomposition dynamics. Across ecosystems, labile organic carbon concentrations consistently decline with increasing soil depth (Adomako et al., 2022, Dragone et al., 2024). Additionally, empirical evidence suggests that limited diffusion fluxes in low-moisture soils can generate or accentuate microscale heterogeneity, thereby increasing habitat diversity within the soil matrix (Dechesne et al., 2008). Consequently, soil heterogeneity is widely recognized as a primary mechanistic determinant underpinning the emergence, stability, and persistence of microbial diversity in terrestrial ecosystems (Nunan et al., 2020).

Although soil microbial communities exhibit extensive phylogenetic diversity, community structure is typically dominated by a limited number of taxa, with the majority occurring at very low relative abundances (Dai et al., 2022). Recent advances in molecular and metagenomic approaches have facilitated more comprehensive investigations of these low-abundance microbial populations, collectively termed the rare biosphere. Despite low abundance, rare taxa can exert disproportionate functional influence on ecosystems. Their genomes often encode unique genetic elements and specialized metabolic pathways that help maintain ecosystem resilience during environmental perturbations. Rare species are increasingly conceptualized as a genetic reservoir, providing an ecological insurance mechanism against environmental change. Following disturbances or episodes of anthropogenic pollution, such taxa are capable of rapid population increases, during which they may stabilize ecosystem functionality by substituting or complementing functions typically carried out by dominant microbial groups. Their metabolic specialization allows them to utilize organic substrates and nutrient pools that remain inaccessible or inefficiently metabolized by dominant taxa (Jousset et al., 2017, Wang et al., 2017). Accordingly, rare microbiota may act as keystone species, directly influencing nutrient turnover and ecosystem functioning. Empirical studies have demonstrated that rare taxa contribute substantially to processes such as organic matter decomposition and nitrogen fixation, in certain contexts exceeding the performance of more abundant taxa in these roles (Wang et al., 2017; Reid and Buckley 2011). Moreover, rare microorganisms can alter the metabolic activities of dominant community members through interspecific interactions, thereby indirectly shaping biogeochemical cycles and ecosystem-level processes (Jousset et al., 2017).

The concept of rarity is intrinsically linked to trophic strategy. The persistence of a stable population within a community can be viewed as the outcome of an organism’s adaptive strategy, whereas environmental fluctuations directly modulate the relative success of distinct trophic modes across spatial and temporal scales (Newton, Shade 2016). Within microbial ecology, two predominant trophic categories are recognized: copiotrophs and oligotrophs. Copiotrophs exhibit rapid growth dynamics in response to resource availability and are specialized for nutrient-enriched environments, while oligotrophs are characteristic of resource-limited ecosystems, where they exploit substrates that are inaccessible or metabolically unfavorable to copiotrophic taxa (Newton, Shade 2016; He et al., 2025). The dichotomy between copiotrophy and oligotrophy shows conceptual convergence with the r–K selection framework, though the latter has been historically applied in macroecological contexts (He et al., 2025). Because microbial ecological studies seldom consider growth kinetics and population stability explicitly within trophic classifications, several authors have advocated for an expanded theoretical framework of microbial life-history strategies that incorporates these additional dimensions of ecological adaptation (Song et al., 2017; Lauro et al., 2009; He et al., 2025).

In numerous studies (Klappenbach et al., 2001; Elser, 2003; Jeyasingh and Weider, 2007; Dethlefsen and Schmidt, 2007), the predominant genomic trait differentiating copiotrophs from oligotrophs has been identified as the copy number of the 16S rRNA operon (rrn). Empirical evidence demonstrates a strong correlation between rrn copy number and the capacity of microorganisms to adjust their growth rates in response to fluctuations in resource availability (Klappenbach et al., 2000). Furthermore, maximum growth rate has been shown to scale positively with rrn copy number, underscoring the ability of copiotrophs to undergo rapid population expansion following nutrient enrichment (Vieira-Silva, Rocha 2010). These genomic and physiological characteristics suggest that the sustained prevalence of copiotrophic populations in nutrient-rich environments is mediated by their capacity to capitalize on episodic nutrient influxes. Importantly, rare or conditionally rare taxa are not necessarily oligotrophic; consistently rare populations are more likely to display oligotrophic strategies, whereas taxa with fluctuating abundance patterns tend to exhibit copiotrophic lifestyles (Lynch, Neufeld 2015; Shade, Gilbert 2015; Newton, Shade 2016).

Comparative genomic analyses further reveal that oligotrophs frequently encode unique genes and specialized metabolic pathways enabling the exploitation of unconventional substrates. They also exhibit elevated efficiency in the degradation of structurally complex organic compounds, though this is often accompanied by reduced intrinsic growth rates. In addition, oligotrophic taxa generally display enhanced stress tolerance and survival under conditions of nutrient scarcity. The integration of these traits is thought to confer ecological advantages in hostile environments and to support the long-term persistence of oligotrophic populations in ecosystems characterized by chronically low fluxes of energy and matter (Ling et al., 2022; He et al., 2025).

Integrating insights from diverse studies, oligotrophic microorganisms can be broadly defined by a suite of genomic and physiological traits indicative of adaptation to nutrient-limited environments. These include reduced genome size, elevated genomic guanine–cytosine content, extended generation times, low intrinsic maximum growth rates, and the capacity to accumulate intracellular storage polymers. Additional hallmarks of oligotrophy encompass high substrate affinity, reduced cell size, greater reliance on passive nutrient transport across the cytoplasmic membrane, a comparatively low surface-to-volume ratio, and ribosomes with enhanced translational fidelity (Schmidt, Konopka 2009; Dragone et al., 2024; Semenov 1991). At the genomic level, oligotrophic taxa are often enriched in genes linked to carbohydrate metabolism, as well as to functional categories associated with virulence, disease, and defense. They also exhibit an increased prevalence of genes encoding pathways for the degradation of structurally complex organic substrates. In contrast, copiotrophic microorganisms generally possess a greater abundance of genetic determinants associated with cell division, cell cycle progression, and sporulation, reflecting their ecological strategy of rapid proliferation under nutrient-enriched conditions (Song et al., 2017).

The current body of evidence for many of the traits associated with oligotrophy remains limited, and the classification of microorganisms as oligotrophic based on single genomic features should be regarded as provisional. Given that oligotrophs are adapted to persist in environments with scarce or intermittent resource availability, it has been hypothesized that oligotrophy may constitute a defining characteristic of uncultivated microorganisms, for which the formulation of suitable growth media and optimal cultivation conditions is often challenging or unattainable. However, empirical studies suggest that oligotrophic lifestyles cannot be robustly inferred from taxonomic affiliation alone. As trophic modes do not represent discrete categories, it is more appropriate to conceptualize them as existing along a continuum, where heterotrophic bacteria from diverse phylogenetic lineages are distributed across a spectrum spanning strongly copiotrophic to strongly oligotrophic life-history strategies (Dragone et al., 2024; Song et al., 2017). The study by Dragone et al., (2024) provided evidence that representative oligotrophic and copiotrophic taxa, broadly associated with soils of low and high carbon content, respectively, can be differentiated within a single dataset. For instance, Pedospheraceae, Ktedonobacteraceae, and unclassified Chloroflexi exhibited strong associations with low-carbon soils, whereas Sphingomonadaceae, Chthoniobacteriaceae, and Solirubrobacteraceae were more frequently detected in carbon-enriched soils. However, comparative analyses across multiple datasets revealed that these associations are not universal, with some exhibiting inverse patterns under different environmental or geographical contexts.

In this study, microbial communities of the total of 31 soil samples, represented by various chernozems, fluvisols, solonets- and solonchak-type soils, were studied. All soil samples were collected in the territory of Specially Protected Natural Areas located in the Rostov Region, Russia. Total soil DNA was extracted and sequenced; metagenomic reads were quality-filtered and used for contig assembly and binning. Quality-filtered bins were analyzed from the perspective of their lifestyle and trophic strategy. The main goal of the study was to understand whether currently recognized oligotrophy markers are suitable for unknown/unculturable microbes delineation and inference of their ecophysiology.

## Materials and methods

### Soil sample collection

Samples were collected during April and May 2022 by envelope method from 0-20 cm depth and stored in aseptic plastic bags under t=4C for no more than a week until further manipulations. A total of 31 samples from a total of 19 unique specifically protected areas were collected: 19 chernozems, 2 chestnut samples (kastanozems), 3 fluvisols, 4 solonetz soils, 2 leptosols and 1 solonchak. Collection dates, coordinates, detailed soil sample types and sampling locations listed in the Supplementary file 1.

### DNA extraction and sequencing

Total soil DNA was extracted using the FastDNA™ Spin Kit for soil. For library preparation, the NEBNext Ultra II DNA Library Prep Kit (NEB) was used according to the manufacturer’s instructions. Genomic DNA was fragmented using the Covaris S220. The quality of the prepared libraries was assessed using the 2100 Bioanalyzer (Agilent) with the DNA High Sensitivity Kit (Agilent). DNA concentration in the preparations was estimated using a Qubit 2.0 instrument. Sequencing was performed on the MGI platform (DNBSEQ) at the Skryabin Institute of Biochemistry and Physiology of Microorganisms, Pushchino, Moscow.

### Quality control of the metagenomic reads

Quality control of the shotgun sequencing data was performed as follows: adapter sequences, low-quality reads (Q < 10), and short reads (< 150 bp) were removed using Trimmomatic v. 0.38 (Bolger, Lohse and Usadel, 2014) with the following specifications: “*SLIDINGWINDOW:4:10 MINLEN:150*”. Low complexity reads and MGI sequencing-specific adapters were removed using fastp program (v. 0.23.4; Chen et al., 2018) with the following options: “*-y -Y 40 –detect_adapter_for_pe --adapter_sequence AAGTCGGAGGCCAAGCGGTCTTAGGAAGACAA --adapter_sequence_r2 AAGTCGGATCGTAGCCATGTCGTTCTGTGAGCCAAGGAGTTG*”. Reads corresponding to human DNA were removed with the minimap2 (v. 2.28; Li, 2018) program using the T2T human genome assembly with the following options: “*minimap2 -t 28 -a -x sr GCA_009914755.4_T2T-CHM13v2.0_genomic.fna*”.

### Contigs and genomes assembly, binning and classification

Quality-controlled reads were assembled into contigs using MEGAHIT (Li et al., 2016) in –meta-large mode with default parameters. Only the contigs with length >1000 b.p. were used for the subsequent analysis. The contigs were binned using two independent binners: Metabat2 (v. 2.12.1, with “*min. contig length 1500”* parameter, the rest are defaults), CONCOCT (v 1.0.0, default mode) (Kang et al., 2019; Alneberg et al., 2014). The bins obtained using these tools were dereplicated and merged into consensus bins using DAS-Tool v. 1.1.5 with *score_threshold* parameter set to 0.4 in order to gather more bins (Sieber et al., 2018). Quality control and assembly statistics of the bins were assessed using the CheckM2 program (Parks et al., 2015; Chklovski et al., 2023). Bins with completeness >90% and contamination <5%, containing 5S, 16S and 23S rRNA and at least 18 tRNA were regarded as high-quality drafts. Bins with completeness >50%, contamination <15%, containing at least one of the 5S, 16S or 23S rRNA genes and minimum 15 tRNA genes were regarded as fine-quality drafts. Bins with completeness and contamination of >50% and <10% respectively were regarded as medium-quality drafts, and as low-quality drafts if below any of the two thresholds. All bins were classified using the GTDB toolkit (‘gtdbtk classify_wf’ module) with the 226-release database (Chaumeil et al., 2020; 2022; Parks et al., 2022).

Raw metagenomic reads were mapped to MAGs contigs and individual genes using coverM (v. 0.7.0; Aroney et al., 2024) with its default minimap2 mapper. Such statistics as trimmed mean coverage, read count, rpkm and tpm were calculated. GCPM (genome copies per million reads) values were calculated based on the MAGs’ contigs coverage using the following formula: 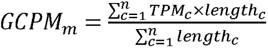, where GCPM_m_ is the GCPM value of MAG m, TPM_c_ is the TPM of contig c in MAG m, length_c_ is the length of contig c in MAG m, and n is the number of contigs in MAG m. Trimmed mean coverage of individual genes were used to calculate generation time of MAGs (see below).

### MAGs generation time prediction

Doubling time for all MAGs was predicted using “gRodon” R package based on the codon usage in ribosomal genes (Vieira-Silva et al., 2010; Okie et al., 2020). Prediction was performed in “metagenome_v2” mode with the “madin” training set. For more robust predictions, data on gene coverage was supplied to gRodon. Gene coverage information was obtained using coverM software as described above.

### MAGs annotation

All genomes were annotated using Prokka default bacterial annotation mode. Carbohydrate-active enzymes were predicted using dbcan3 (v.5.2.1; Zheng et al., 2023) default settings. Several features were considered in the scope of oligotrophy/copiotrophy delineation (Table 1).

**Table 1.** Microbial features considered as oligotrophy markers or related to oligotrophy lifestyle.

| Feature | Characteristics in oligotrophs | References |
| --- | --- | --- |
| Genome size | Typically, smaller | Dragone et al., 2024 |
| Copies of rRNA operons | Typically, low copy number | Klappenbach et al., 2001; Elser, 2003; Jeyasingh and Weider, 2007; Dethlefsen and Schmidt, 2007 |
| Generation time | Typically, >5 hours of cell doubling time | Vieira-Silva et al., 2010; Okie et al., 2020 |
| Chemotaxis proteins | Typically, low copy number or absent | Lauro et al., 2009; Clerc et al., 2022; Barnett et al., 2023 |
| Two-component system (2CS) protein abundance | Typically, low copy number | Noell et al., 2023 |
| C1-substrate degradation | Often a major trophic strategy | Peres et al., 2023; Halsey, Carter and Giovannoni, 2012; Schmider et al., 2024 |
| Dissipation of low-molecular weight carbon | Often a major trophic strategy | Ghuneim et al., 2021; Berggren et al., 2010 |
| Complex biopolymers degradation (i.e. CAZymes) | Less present or absent | Leu et al., 2022; López et al, 2023; |
|  |  | Sun et al., 2023 |
| Ability to oxidize molecular hydrogen | Often present | Lappan et al., 2023; Lau et al., 2016 |

**Table 2.** summarizes data on the metabolic pathways that were identified in more than 50% of phylum genomes.

| Phylum | What substrates can be utilized according to genes in >50% of phylum genomes |
| --- | --- |
| Acidobacteriota (N=40) | benzoyl-coa, carbon monoxide, formate, OS, carbohydrates |
| Actinomycetota (N=32) | benzoyl-coa, carbon monoxide, formate, OS |
| Bacteroidota (N=37) | benzoyl-coa, carbohydrates |
| Bacteroidota_A (N=8) | benzoyl-coa, formate, carbohydrates |
| Chloroflexota (N=6) | benzoyl-coa, carbon monoxide, formate, OS, carbohydrates |
| Desulfobacterota E (N=1) | benzoyl-coa, formate |
| Desulfobacterota B (Binatota) (N=12) | benzoyl-coa, carbon monoxide, formate, OS |
| Gemmatimonadota (N=12) | benzoyl-coa, formate, methanol |
| Methylomirabilota (N=3) | benzoyl-coa, formate, carbon monoxide, OS |
| Myxococcota (N=9) | benzoyl-coa, formate, OS, carbohydrates |
| Planctomycetota (N=6) | benzoyl-coa, carbohydrates |
| Pseudomonadota (N=50) | benzoyl-coa, methanol, OS, AMO/PMO, carbohydrates, proteins and peptides |
| Verrucomicrobiota (N=23) | benzoyl-coa, formate, OS, carbohydrates |
| Tectomicrobia (N=1) | benzoyl-coa, formate, carbon monoxide, hydrogenase, methanol, OS |
| Patescibacteriota(N=3) | benzoyl-coa |
| Krumholzibacteriota (N=1) | benzoyl-coa |
| Eisenbacteria (N=1) | benzoyl-coa, proteins and peptides |

In this study, genome size was assessed by checkm2, copies of rRNA operons - by barrnap, generation time - by gRodon R package, and functional capabilities were inferred from Prokka annotation results. The C1-substrate degradation feature was addressed by analyzing the presence and the quantity of methanol-, methane-, formate-and CO-oxidizing enzymes. Ability to oxidize H2 was inferred from the presence and the quantity of NiFe-hydrogenases. For the dissipation of low-molecular weight carbon, benzoyl-CoA utilization pathways and organosulfonate utilization pathways were analyzed.

### Statistics

Principal component analysis (PCA) was performed using prcomp(center = TRUE, scale. = TRUE) command in R. The optimal number of clusters was evaluated using clusGap(pc_scores, FUN = kmeans, K.max = 10, B = 50) and silhouette() commands from the “cluster” library. Significance of genomes clustering on PCA plot was done using kruskal.test() plus post hoc pairwise.wilcox.test() with Benjamini-Hochberg adjustment. Cluster stability assessment was done using clusterboot(B = 100, bootmethod = "boot",clustermethod = kmeansCBI,k = 6) function from “fpc” library.

## Results and discussion

### Assembly and binning of metagenomic reads

After quality control, metagenomic samples were characterized by a varying number of reads. The smallest sample comprised 7953643 reads, the largest – 99558815, while average sample size was ∼50.7M reads. MEGAHIT assembly of 31 samples resulted in a total of 5879100 contigs after discarding <1000 b.p. contigs; smallest number of which was in sample 1702 – 5884, and the largest – 535111 in sample 1819. The largest contig of 585712 b..p. was assembled from sample 1701. Mean coverage of contigs per sample ranged from 2.5 to 7.5x. Using the assembly and binning methods described above, a total of 434 bins were formed. After discarding all bins that fall under quality metrics described above and all bins for which it wasn’t possible to estimate any of the Table 1 criteria, 249 bins left. Maximum completeness level of 100% and minimum contamination level of 0% was obtained for two Bacteroidota bins: D_102 and D_124. Maximum genome size of 11.6 mb was observed for Myxococcota (c Polyangia) MAG 1603_0 (completeness and contamination are 97.2 and 4.3% respectively). Minimum genome size was observed for two Patescibacteriota (c Saccharimonadia) MAGs: 0.58 mb for D_143_2 and 0.61 mb for D_106 (with completeness levels of 90 and 77% respectively, estimated by checkm2 for small-genome bacteria). The mean contamination value for all genomes was 5.1%. The average genome size was 4,104,404 bp.

The average genome assembly completeness for all genomes was 79%. Mean completeness values for individual phyla were as follows: Acidobacteriota – 79%, Actinomycetota – 73%, Bacteroidota – 85%, Bacteroidota A – 80%, Chloroflexota – 70%, Desulfobacterota E – 51%, Desulfobacterota B (Binatota) – 76%, Gemmatimonadota – 79%, Methylomirabilota – 75%, Myxococcota – 82%, Planctomycetota – 70%, Pseudomonadota – 81%, and Verrucomicrobiota – 81%. Single genomes showed completeness values of: Tectomicrobia – 65%, Patescibacteriota – 62%, Krumholzibacteriota – 52%, and Eisenbacteria – 76%. All further analyses were done with this 246 MAGs dataset. A detailed table with all taxonomy, statistics, abundance and other information is presented in the Supplementary file 2.

### Generation time prediction results

The slowest cell doubling rate was observed in an Actinomycetota genome (c UBA4738, o UBA4738, f HRBIN12, g DASSJM01) at 15.6 hours. On average, the longest generation times were found in Methylomirabilota (11.2 h) and Actinomycetota (8.9 h). Representatives of Gemmatimonadota (8.1 h), Desulfobacterota E (7.7 h), and the single Eisenbacteria genome (8.3 h) also had prolonged division times. The average doubling time for representatives of Pseudomonadota (4.8 h), Verrucomicrobiota (5 h), Acidobacteriota (5.9 h), Chloroflexota (5.8 h), Myxococcota (6.1 h), Bacteroidota A (5.6 h), Planctomycetota (7.1 h), Desulfobacterota B (Binatota) (5.6 h), and for the single genomes of Patescibacteriota (5.5 h), Krumholzibacteriota (5.8 h), and Bacteroidota (4.5 h) ranged from 5 to 7 hours. The single Tectomicrobia genome had a generation time of 3.5 hours. The highest doubling rates were observed in certain Pseudomonadota representatives (∼0.6 hours).

Summary on genome statistics and some key annotation results is presented in Figure 1.

**Figure 1.**
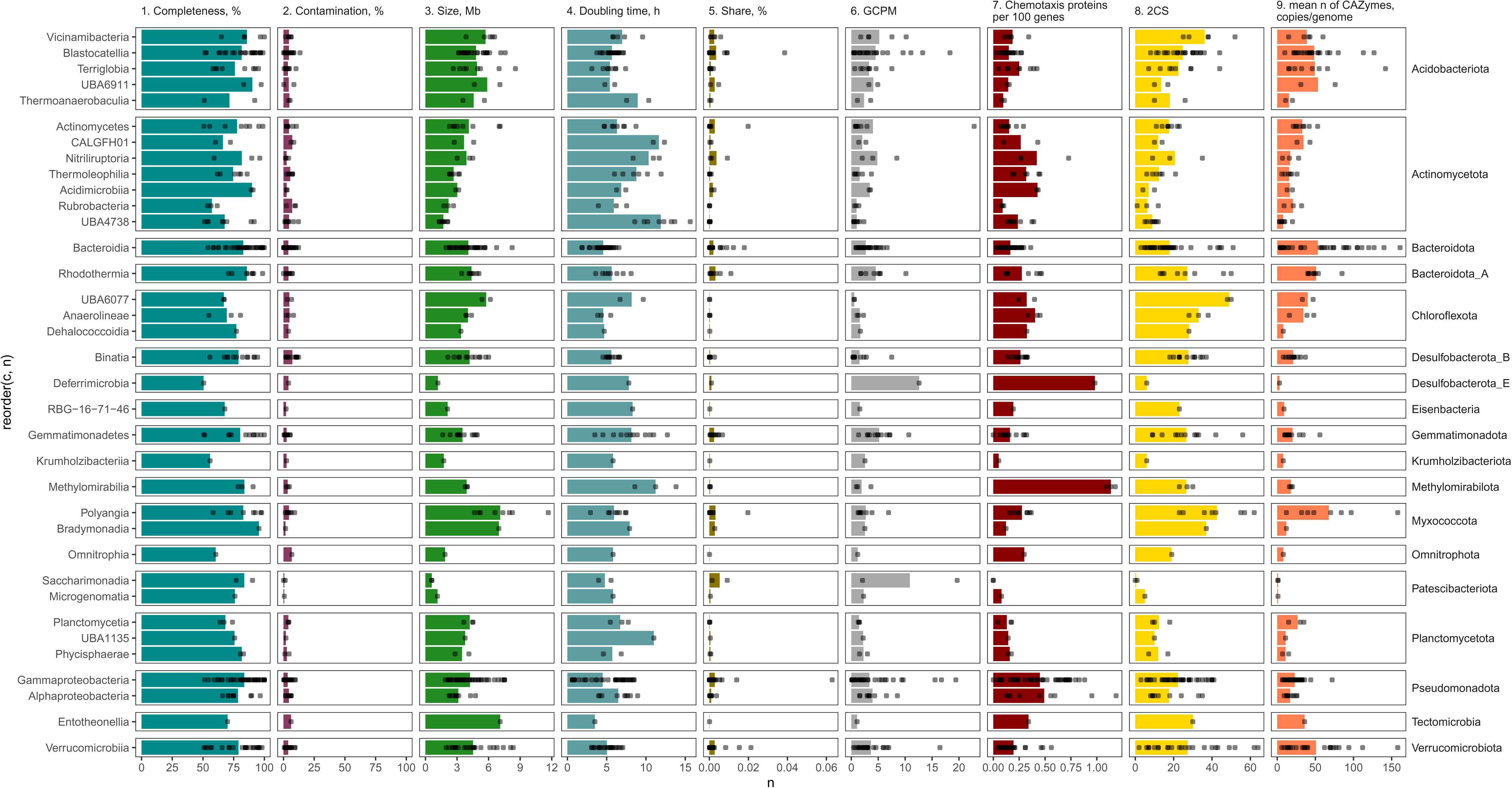
Multi-panel barplot comparing genomic and ecological traits across multiple bacterial phyla and classes. Each dot represents an individual MAG, and MAGs are grouped by taxonomic classification on the y-axis. The figure displays nine panels, each quantifying a different genomic or predicted phenotypic trait. The first two panels show genome quality metrics: completeness ranges from 0–100% and contamination ranges from 0–100%. The third panel shows genome size spanning 0–12 megabases, while the fourth panel displays predicted doubling time from 0–15 hours. The fifth panel indicates relative community abundance on a scale of 0–0.06%. The sixth panel shows GCPM (genome copies per million of metagenomic reads, used as a normalized abundance metric) ranging from 0–20. The seventh panel presents chemotaxis protein frequency per 100 genes from 0–1. The eighth panel shows two-component system (2CS) gene counts from 0–60. Finally, the ninth panel displays the mean number of carbohydrate-active enzyme genes per genome, ranging from 0–150 copies.

**Figure 2.**
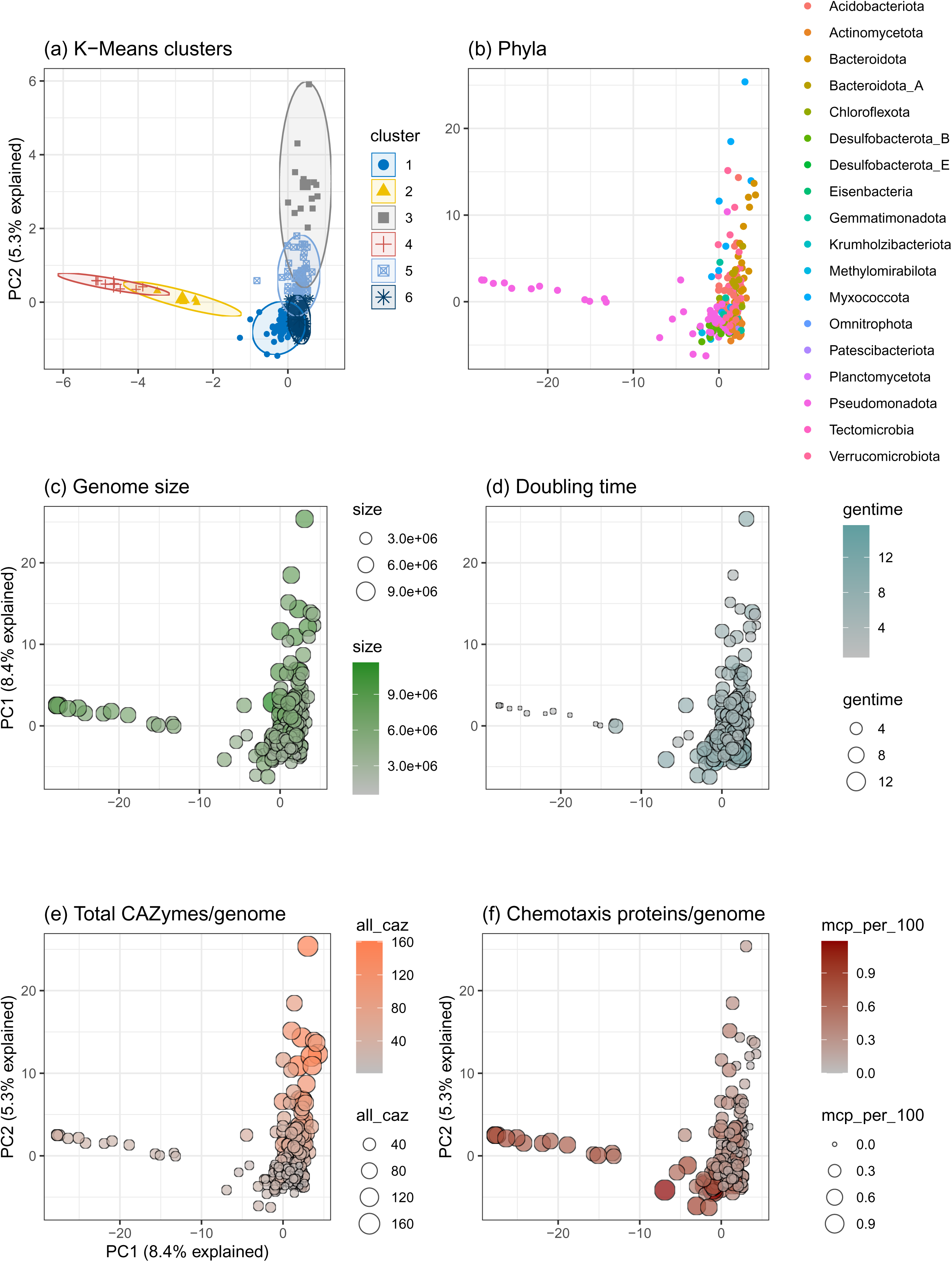
Multiple-facets Principal Component Analysis (PCA) biplot displaying PCA results with 6 clusters (a), mapped phylum (b), genome size (c), doubling time (d), total CAZymes and Chemotaxis proteins share (e) assignments to each genome (represented by dots).

The largest genomes (6–12 Mb) are predominantly found in Myxococcota (particularly Polyangia and Bradymonadia), Planctomycetota, and certain Acidobacteriota classes. In contrast, the smallest genomes (<3 Mb) cluster in Patescibacteria (Microgenomatia, Saccharimonadia), some Chloroflexota lineages, and candidate phyla. The predicted generation time shows pronounced variation across taxa, with Gammaproteobacteria and Bacteroidia exhibiting rapid doubling times (0–5 hours), consistent with their known copiotrophic ecology in high-nutrient environments. In stark contrast, Acidobacteriota (especially Terriglobia and Blastocatellia), Chloroflexota, and Methylomirabilota display substantially longer doubling times (10–15 hours or longer), reflecting their adaptation to resource-limited conditions or specialized niche exploitation. Myxococcota, Planctomycetota, and Pseudomonadota carry 40–60 copies of 2CS genes, reflecting their investment in complex regulatory networks for coordinating group behaviors (in Myxococcota), detecting nutrients and competitors, and executing coordinated physiological responses. In contrast, Patescibacteria, some Chloroflexota, and candidate phyla carry fewer than 10 2CS genes, indicating minimal regulatory complexity. This reduction in two-component systems is consistent with obligate symbiosis (where a host handles environmental sensing) or extreme specialization where a streamlined regulatory architecture suffices. The 2CS counts validate earlier findings that regulatory complexity is a hallmark of copiotrophic bacteria capable of sensing and responding dynamically to environmental fluctuations, whereas oligotrophs or strict specialists have reduced regulatory overhead. Chemotaxis protein frequency per 100 genes shows distinct phylogenetic clustering. Taxa such as Myxococcota (characterized by social motility and coordinated predation) and Pseudomonadota consistently carry high chemotaxis gene densities (0.5–1.0 per 100 genes). These motile, competitive copiotrophs invest in chemotaxis machinery to actively seek nutrients and engage in metabolic competition. Conversely, Patescibacteria, non-motile Chloroflexota classes (Dehalococcoidia, Anaerolineae), and several Acidobacteriota show minimal or zero chemotaxis genes. Bacteroidota, Planctomycetota, and Verrucomicrobiota carry 80–150+ CAZyme genes, positioning them as specialist polymer degraders capable of attacking cellulose, chitin, pectin, xylan, and other complex structural polysaccharides. These high CAZyme counts enable rapid colonization and degradation of plant biomass and algal detritus, a characteristic copiotrophic strategy in biomass-rich environments. Conversely, Patescibacteria, Methylomirabilota, and some Chloroflexota carry fewer than 20 CAZyme genes on average, reflecting reliance on low-molecular-weight substrates (amino acids, sugars), C1 compounds (methanol, methane, formate), or host-derived nutrients rather than independently degrading complex polymers. Copiotrophs - represented predominantly by Gammaproteobacteria, Bacteroidia, Myxococcota, and Planctomycetota - consistently display large genomes (6–12 Mb), rapid growth (low doubling times), extensive two-component signaling systems (40–60 copies), frequent chemotaxis genes, and high CAZyme inventories (80–150+ genes). These traits enable rapid response to nutrient influxes, active resource competition, and efficient degradation of complex polymers. At the opposite extreme, oligotrophs and extreme specialists - Patescibacteria, Chloroflexota (particularly Dehalococcoidia), and Methylomirabilota - maintain small, streamlined genomes (<3 Mb), slow generation times, minimal regulatory complexity (few 2CS genes), limited or absent chemotaxis, and low CAZyme counts. These genomes reflect adaptation to chronically resource-limited conditions, fixed ecological niches, or specialized metabolic dependence on C1 compounds and hydrogen. Intermediate taxa, including various Acidobacteriota, Verrucomicrobiota, and Actinomycetota classes, exhibit intermediate trait values and variable strategies, suggesting functional diversity within these groups that may reflect niche partitioning or heterogeneous selective pressures across soil microhabitats.

Three Methylomirabilota members (MAGs 1707_11, 1801_60, 1805_17) here provide a case of partial oligotrophic markers: they are slowDgrowing (predicted doubling time exceeds 10 hours), nicheDspecialized C1 oxidizers adapted to lowDsubstrate environments, however, the studied representatives are regulationDrich and motile, optimized for sensing C1-electron donors and nitrogen-containing electron acceptors (Versantvoort et al., 2018; Nie et al., 2023).

Krumholzibacteriota MAG 1805_30 shows moderate to high genome size, slow doubling time (reflecting oligotrophic specialization on low-flux aromatic substrates), moderate two-component systems and chemotaxis genes (for locating aromatic hotspots in stratified environments), and low to moderate CAZyme counts (reflecting specialization on aromatics rather than polysaccharides). It has low regulation and chemotaxis potential and low CAZyme abundance. Moreover, the genome encodes benzoyl-CoA reductase and other proteins related to monoaromatic degradation through benzoyl-CoA. Together with low abundance metrics in the community, this all points to the clear oligotrophic strategy of a rare biosphere member.

Eisenbacteria is one of the most poorly characterized phyla now and its members are regarded as extremely rare biosphere microbes (White III et al., 2023). Here, Eisenbacteria MAG 1818_17 (c RBG-16-71-46, g WS-7), exhibited relatively small genome size, moderate doubling time, 2CS and chemotaxis proteins abundance, and low CAZyme counts - consistent with an oligotrophic, specialist lifestyle. Their rarity and lack of cultivation suggest they occupy highly specialized niches with limited overlap with better-studied phyla. In Zhong et al., (2022), Eisenbacteria MAG were characterized by nutrient-attracted motility and trophic strategy oriented towards proteins and peptides. Data obtained here agrees with that, since we also observe nutrient-sensing potential and protein-related metabolism (MAG 1818_17 encodes different endopeptidases and proteases). Eisenbacteria may represent a case of specialized, extremely rare oligotrophs, still having other, not purely oligotrophic, characteristics.

### Annotation results

Many representatives of the presented phyla were found to possess genes for aerobic, anaerobic, and hybrid pathways of benzoate and derivative transformations. Genes of the benzoyl-CoA pathway are thought to be associated with the degradation of monoaromatic compounds of both plant and industrial origin. Aromatic compounds derived from lignin, flavonoids, quinones, aromatic amino acids, or fossil fuels are recognized as some of the most widespread classes of organic molecules in nature. However, they are often considered recalcitrant in the environment due to the high chemical stability of the aromatic ring (Porter, Young, 2014). Therefore, the ability to metabolize such compounds may play a significant role in the ecology of rare microbiota, providing the capacity to use substrates that are inaccessible to many bacteria. The initial mechanisms for degrading these compounds are diverse, but they converge on the formation of a common intermediate - benzoyl-CoA.

Genes of the benzoyl-CoA pathway were most abundant in representatives of the Binatota (especially with high copy numbers of 4-hydroxybenzoyl-CoA reductase in MAGs 1816_22, 1502_3, and 1815_8) and Acidobacteriota (notably in MAGs 1810_1 and 1816_97 from c. Vicinamibacteria). All representatives of Pseudomonadota, Methylomirabilota, Chloroflexota, Verrucomicrobiota, Planctomycetota, Myxococcota, Bacteroidota, and most Gemmatimonadota, Actinomycetota, and Bacteroidota A possessed at least one gene encoding enzymes of this pathway. The same was true for a few detected representatives of Desulfobacterota_E, Eisenbacteria, Krumholzibacteriota, Tectomicrobia, and Patescibacteriota.

Other enzymes implicated in the decomposition of complex organic substrates include those involved in coniferyl alcohol transformation, given that coniferyl alcohol is recognized as a major lignin precursor and may be associated with lignin breakdown products (Buraimoh et al., 2017; Nishimura, 2013). These enzymes were identified in a single representative of Myxococcota and in approximately half of the metagenome-assembled genomes (MAGs) belonging to Pseudomonadota. Notably, MAG D_40_2 from Alphaproteobacteria encoded two enzymes of this pathway, while the greatest number of MAGs harboring a single enzyme were classified within the genus Pseudomonas_E.

From another perspective, the final metabolic products of complex organic molecule degradation may themselves be considered recalcitrant, as their further metabolism is limited to microorganisms possessing highly specific enzymatic systems.

Methanol is produced in significant quantities during mineralization processes in nature, mainly as a result of the breakdown of methyl esters present in plant-derived components such as pectin and lignin (Hektor, Kloosterman, Dijkhuizen, 2000). Among Pseudomonadota and Gemmatimonadota, over half of the detected representatives possessed genes encoding at least one of the three methanol-metabolizing enzymes. In Verrucomicrobiota, fewer than half had these enzymes. Nearly half of Acidobacteriota representatives contained genes involved in methanol conversion. Of the two identified genera within Methylomirabilota, only one representative harbored both NAD-dependent methanol dehydrogenase and methanol dehydrogenase activator. In Actinomycetota, Bacteroidota, and Bacteroidota A, the enzyme was present in less than 40% of representatives. In the relatively few Planctomycetota and Tectomicrobia, NAD-dependent methanol dehydrogenase was found in only one representative, whereas about half of Chloroflexota and Myxococcota possessed this enzyme.

Formate is another common product of bacterial fermentation of larger and more complex carbon sources. It accumulates in environments lacking terminal electron acceptors and can serve as a substrate for subsequent metabolic processes such as acetogenesis, methanogenesis, and hydrogen production; thus, formate plays a key role in anaerobic syntrophic associations (Bar-Even, 2016). It is believed that standalone formate conversion does not occur in pure cultures due to thermodynamic constraints (Lim et al., 2012). Genes associated with both aerobic and anaerobic formate metabolism were generally widespread in our dataset. The highest prevalence was found among Actinomycetota, Acidobacteriota, and Binatota. At least half of the representatives of Bacteroidota A, Chloroflexota, Gemmatimonadota, Methylomirabilota, Myxococcota, Verrucomicrobiota, Desulfobacterota E, and Tectomicrobia also encoded genes for formate metabolism.

One representative of Actinomycetota carried genes for methane monooxygenase (MAG 1812_47) and ammonia monooxygenase (MAG 1705_9), and over half of Pseudomonadota MAGs encoded ammonia monooxygenase. These enzymes catalyze the initial transformation of ammonia (AMO) and methane (pMMO), representing the first and often rate-limiting step in their respective metabolic pathways. Ammonia and methane, in turn, are key intermediates in biological cycles.

From a different perspective, a competitive adaptation may be the ability to efficiently utilize compounds present in persistently low concentrations in the environment. [NiFe]/[NiFeSe]-hydrogenases are enzymes catalyzing molecular hydrogen (H2) oxidation. It has been demonstrated that some aerobic soil actinobacteria and acidobacteria are capable of survival by scavenging H2 from atmospheric trace concentrations. Notably, hydrogen oxidation activity increases under carbon limitation. It has been proposed that soil microorganisms possessing high-affinity hydrogenases may be especially competitive, enabling them to exploit a dependable energy source under unstable conditions (Greening C. et al., 2014; Greening C. et al., 2016). These genes were most prevalent among Actinomycetota MAGs. Additionally, at least one subunit gene of periplasmic [NiFe]/[NiFeSe] hydrogenase was detected in some representatives of Methylomirabilota, Gemmatimonadota, Planctomycetota, Bacteroidota, Myxococcota, and Acidobacteriota.

Our metagenomic analysis also revealed another hydrogenase associated with metabolism of gases that are consistently present at low concentrations in the atmosphere - CO. The enzyme carbon monoxide dehydrogenase/acetyl-CoA synthase (CODH/ACS) catalyzes reactions converting CO2 to CO and CO2 to a methyl group, which then enters acetyl-CoA, but in certain bacteria and archaea, CODH/ACS can also operate in reverse. Traditionally, the reductive acetyl-CoA pathway is considered characteristic of methanogens and acetogens; however, the presence of CODH/ACS homologs has also been reported in numerous uncultured anaerobic organisms (Adam, Borrel, Gribaldo, 2018).

In a 2019 study (Cordero P. R. F. et al., 2019), it was demonstrated that atmospheric CO oxidation supports bacterial survival under nutrient limitation. Furthermore, CO has been shown to play an important role in the persistence of microbial communities in severely oligotrophic environments, such as polar deserts (Ji et al., 2017) and volcanic deposits (King, 2003). Genes encoding carbon monoxide dehydrogenase were present in all assembled genomes of Methylomirabilota and Binatota. The largest numbers of gene copies encoding the three subunits of this enzyme were observed in MAG 1805_17 (Methylomirabilota), 1603_12 and 1816_22 (Binatota), and 1501_27 (Chloroflexota). More than half of the representatives of Pseudomonadota, Acidobacteriota, and Actinomycetota possessed genes encoding this enzyme, as did the single detected representative of Tectomicrobia.

Some microorganisms can degrade complex compounds not only under severe nutrient limitation but also under conditions where essential metabolic substrates are scarce. One example is the acquisition of sulfur from organosulfonate compounds during sulfur depletion. In most bacteria, this system comprises enzymes that catalyze desulfonation of various groups, the oxidation of organosulfonate compounds, transport proteins mediating uptake from the environment, and enzymes involved in dissimilation of different sulfur compounds (Ellis, 2011).

Among Binatota, genes encoding dimethylsulfide dehydrogenase - an enzyme catalyzing the oxidation of dimethylsulfide (DMS) to dimethylsulfoxide, a volatile organic compound (VOC) - were also detected. Traditionally, DMS and related compounds are considered to be primarily associated with marine ecosystems; however, mounting evidence points to their potential ecological relevance in terrestrial habitats as well. The production of VOCs in terrestrial environments may be linked to increased plant resistance to diverse abiotic stresses (Rinnan et al., 2014). DMS has been observed to form in peat bogs and during the decomposition of plant residues in soil (particularly crucifers), and there are reports of DMS emissions by maize, alfalfa, and wheat (Schäfer, Myronova, Boden, 2010). DMS is the product of microbial degradation of DMSP (Bentley, Chasteen, 2004), which in turn can be synthesized by certain plants, such as a few Spartina species, some sugarcanes, and the coastal strand plant *Wollastonia biflora* (Otte M. L. et al., 2004).

Several authors have reported that DMSP can be generated as a result of industrial processes involving organic matter transformation (e.g., wood pulp production, petroleum processing, manure management, and wastewater treatment) (McDevitt C. A. et al., 2002), and more broadly in the food and brewing industries, as well as in agriculture and livestock farming (Schäfer, Myronova, Boden, 2010). Another DMSP transformation pathway is demethylation via a specific enzyme, resulting in methanethiolpropionate, and subsequently acetaldehyde (which is assimilated as a carbon source) and the reactive gas methanethiol (MeSH), which can be utilized as a sulfur source (Li et al., 2024). Dimethylsulfoniopropionate demethylases were detected in almost half of Actinomycetota representatives and in some of Pseudomonadota and Acidobacteriota.

Dimethyl sulfone (DMSO2) may also serve as an alternative sulfur source, supporting bacterial survival in low-sulfate environments (Gonzalez et al., 2024), as it too can be metabolically converted for carbon and electron acquisition (Soule et al., 2020). The enzyme responsible for DMSO2 conversion - dimethylsulfone monooxygenase - was detected in certain representatives of Acidobacteriota, Actinomycetota (especially the genus g DASSJM01), Binatota, Chloroflexota, Gemmatimonadota, Methylomirabilia, and Pseudomonadota.

Organosulfonate metabolism was a widespread feature rather than specialized microbial metabolism and was found in 12/18 phyla. However, substrate preferences were observed at the phylum level. The most diverse and abundant were genes encoding for aliphatic sulfonate transport and binding proteins. Desulfonating enzymes, such as alkanesulfonate monooxygenase, taurine dioxygenase and sulfoacetaldehyde acetyltransferase were found across Acidobacteriota, Actinomycetota, Binatota, Pseudomonadota, Verrucomicrobiota, Methylomirabilota and Chloroflexota members in more >2 genomes. Taurine dioxygenase occurrence was mostly restricted to gammaproteobacteria. Metabolic potential for sulfolactate and methanesulfonate utilization was partially found in Pseudomonadota and Acidobacteriota MAGs.

### Exploratory analysis and grouping of genomes based on the oligotrophy markers

To compress multidimensional data into meaningful 2D representation, PCA analysis using multiple oligotrophy markers was performed. Initially, multiple runs of PCA tests were performed with different types, quantity and level of grouping of all selected variables in order to find such a combination that splits genome datasets into meaningful, statistically significant clusters. It turned out that including all proteins individually provides the most resolved picture. In the end, a dataset of 347 variables (different genome metrics and protein copies) and 246 observations (genomes) was used for the final PCA run. According to the clusGap() analysis, the optimal number of clusters was 6. This was confirmed by the Kruskall-Wallis test of differences between clusters along the first two PCs, following the pairwise Wilcoxon test with Benjamini-Hochberg p.value adjustment. Only two pairwise comparisons (clusters 5 and 6 along the PC1 and clusters 4 and 5 along the PC2) had p-values over the 0.05 threshold. Average cluster width according to silhouette analysis was 0.47, and bootstrap analysis on cluster stability showed 0.86, 0.42, 0.84, 0.63, 0.65, 0.79 for clusters 1 to 6 respectively, which are moderately good clustering results.

Variables considered for PCA analysis have shown good mapping on the final plot, again strengthening the assumption that oligotrophy markers can group genomes into reasonable clusters (Figure 3).

**Figure 3.**
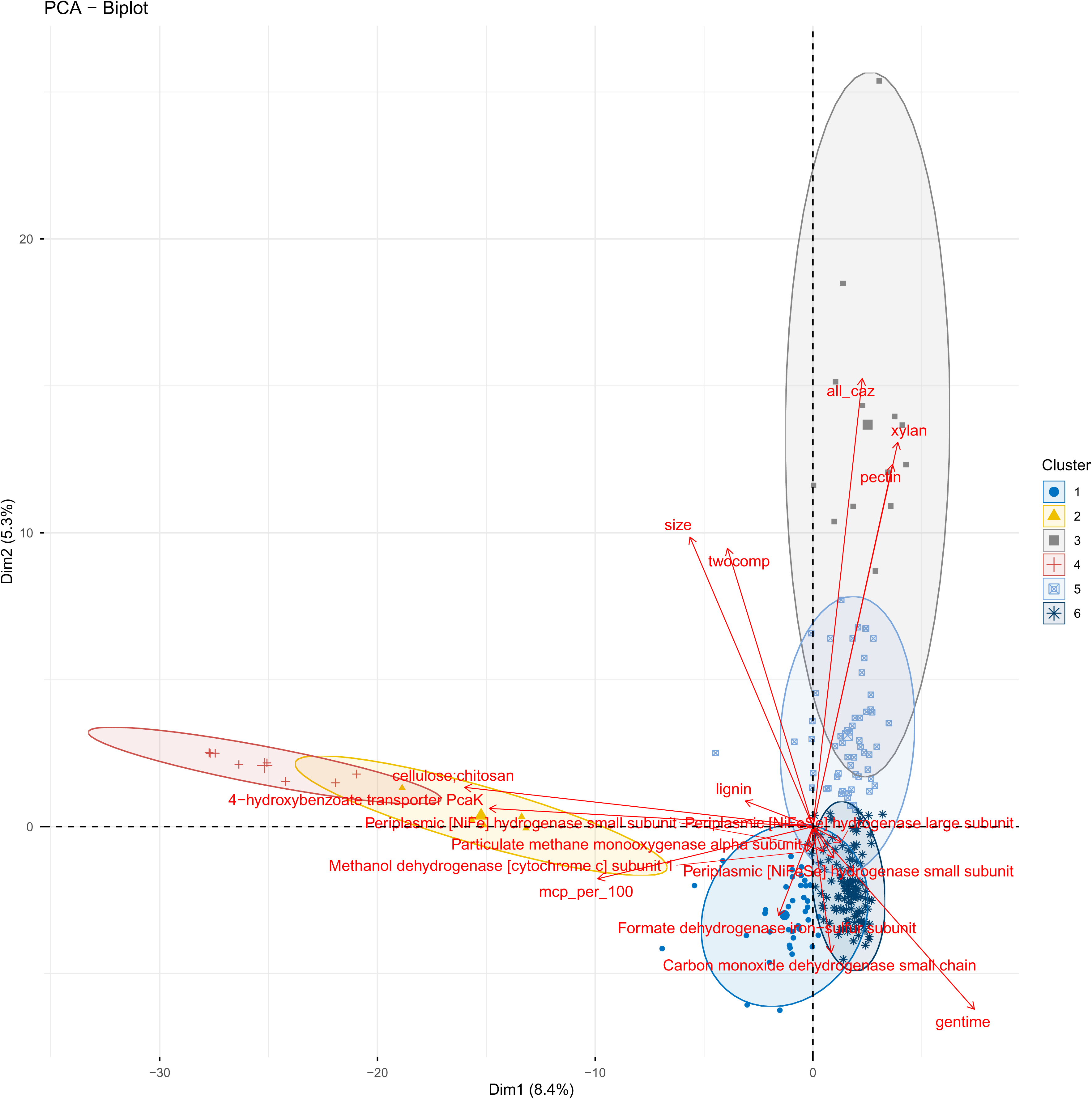
Principal Component Analysis (PCA) biplot displaying relationships between bacterial genomic and functional features across six distinct clusters.

Figure 3 reveals a complex landscape of bacterial ecological strategies, with clear differentiation between clusters that likely represent distinct life-history strategies and metabolic capabilities of the studied genomes.

Two-component systems and chemotaxis proteins abundance as markers for oligotrophs appear validated in this dataset. The biplot shows twocomp and mcp_per_100 as distinct variables in the PCA space, indicating they vary significantly across bacterial samples and contribute meaningfully to the clustering pattern. This suggests that: clusters with high twocomp and mcp_per_100 likely represent fast-growing copiotrophs with rapid environmental response capabilities and active motility. Clusters with low values may represent oligotrophic or specialized bacteria relying on passive nutrient uptake. The prominence of hydrogenases (NiFe, NiFeSe) and methane/methanol appear to be important in clustering as well. These metabolic pathways are typically associated with specialized, slow-growing oligotrophs adapted to nutrient-poor environments or specific redox niches. The presence of carbohydrate-active enzymes (CAZymes) targeting cellulose, chitosan, xylan, pectin, and lignin indicates that the dataset includes decomposers and primary colonizers of plant biomass. The all_caz (total sum of CAZymes per genome) variable shows strong positive association with certain clusters, separating: The size variable in the biplot links to the broader ecological pattern. Larger genomes typically correlate with metabolic versatility and are associated with copiotrophs or facultative specialists - organisms with flexible lifestyles.

The six clusters likely represent: 1) Clusters 4, 2 and 3: Fast-growing generalists (possibly high twocomp, mcp, CAZymes, genome size) - copiotrophs capable of rapid response to nutrient influxes; 2) Part of the cluster 6: Hydrogenotrophs (high hydrogenase genes, possibly lower CAZymes, distinct metabolic profile) - specialized oligotrophs using HD as energy source; 3) Part of the clusters 1 and 6: Methanotrophs/methylotrophs (the presence of pMMO/formate dehydrogenase/methanol dehydrogenase/CO dehydrogenase, distinct energy metabolism) - specialized bacteria oxidizing C1 compounds; 4) Clusters 5 and 3 members: Polysaccharide degraders (high CAZyme diversity for specific polymers, moderate growth capacity) - decomposers with semi-specialized lifestyles; 5) Aromatic degraders, distributed among all clusters, but late stage degrades are concentrated in 1 and 6 clusters. Bacteria adapted to phenolic/aromatic niches; they encode Benzoyl-CoA reductase, Cyclohexadiencarbonyl-CoA hydratase and other enzymes finalizing monoaromatics degradation; 6) Cluster 6: minimal genome specialists (potentially small genome size, low regulatory complexity, specialized metabolism) - highly adapted oligotrophs.

Crucially, the first two components explained no more than 15% of initial data variance. Clusters 1, 5 and 6, which represent different oligotrophic groups, are highly similar to each other and clustered tightly. The relatively low variance explained by the first two dimensions suggests that bacterial variation is distributed across many features - typical for multi-faceted genomic datasets. This also suggests that more variables, did not considered here, should be included for the proper and more resolved delineation of different oligotrophs groups, likely the variables that reflect more detailed substrate preferability or microbe-microbe interactions. Still, this analysis clearly validates the use of genome size, generation time, 2CS and chemotaxis protein frequencies combined as ecological markers. The fact that 2CS and chemotaxis proteins appear as meaningful variables in the PCA biplot, contributing to cluster separation, demonstrates they are informative for distinguishing bacterial lifestyle strategies. Clusters with high values likely represent organisms with copiotrophic traits (rapid sensing, motility, competitive advantage), while clusters with low values represent oligotrophic or specialized strategies where such regulatory complexity is unnecessary.

## Conclusions

In this study, 246 bacterial genomes, assembled from different type soil metagenomes, were studied from the perspective of their trophic strategy and ecophysiology. Several genomic and functional traits were analyzed as predictors for oligotrophy/copiotrophy lifestyle. It has been shown that genome size, doubling time, complexity of carbon sources preferred and chemotaxis and signaling flexibility successfully cluster bacterial genomes into at least 6 clusters. Doubling time, abundance and diversity of carbohydrate-activate enzymes and chemotaxis proteins turned out as one of the most effective predictors. Oligotrophy as a life strategy was inferred for Methylomirabilota, Krumholzibacteriota, and Eisenbacteria MAGs. Copiotrophy as a life strategy was inferred for the majority of Myxococcota, Bacteroidota, Gammaproteobacteria and Verrucomicrobiota MAGs.

## Supporting information

Collection dates, coordinates, detailed soil sample types and sampling locations listed in the Supplementary file 1.

A detailed table with all taxonomy, statistics, abundance and other information is presented in the Supplementary file 2.

## Credit author statement

K.Y.N. performed DNA extraction, library preparation and sequencing; D.K.A. formulated initial hypotheses, performed metagenomic binning and analysis and wrote the final draft; S.V.V. wrote the first draft and assisted with manuscript writing, editing and preparation for submission; K.K.I., M.P.A. and S.V.V. performed annotation and in-depth genome analysis; M.T.M. and P.E.V. acquired funding, supervised the project, contributed to manuscript writing, reviewed and revised the final draft.

## Data availability

Currently, all sequencing data are available upon request.

